# CD4 binding-site antibodies induced by a subtype B HIV-1 envelope trimer

**DOI:** 10.1101/2022.03.23.485469

**Authors:** Anna Schorcht, Tom L.G.M. van den Kerkhof, Jon Torres, Edith Schermer, Celia C. LaBranche, Ilja Bontjer, Mitch Brinkkemper, Naveed Gulzar, Alvin X. Han, Judith Burger, Gabriel Ozorowski, Jamie K. Scott, Hanneke Schuitemaker, David Montefiori, Marit J. van Gils, Andrew B. Ward, Rogier Sanders

## Abstract

In the last decade considerable advances have been made towards the design of HIV-1 vaccines that induce neutralizing antibodies (NAbs). Despite the progress, no vaccine is able to consistently elicited broadly neutralizing antibodies (bNAbs). Here we present a case study of a rabbit that was immunized with a subtype B native like envelope glycoprotein (Env) trimer, AMC016 SOSIP.v4.2, with a dense and intact glycan shield, followed by a trivalent combination of subtype B trimers. After the priming phase serum from this animal neutralized several heterologous subtype B neutralization resistant (tier 2) viruses. Subsequent immunization with the trivalent combination of subtype B trimers further increased the breadth and potency of the NAb response. EM based polyclonal epitope mapping revealed that a cross reactive CD4 binding-site (CD4bs) antibody response, that was present after priming with the monovalent trimer and boosting with the trivalent combination, was most likely responsible for the broad neutralization. While anecdotal, this study provides proof-of-concept that native-like Env trimers are capable of inducing CD4bs-directed bNAb responses and should guide efforts to improve the consistency with which such responses are generated.

## Introduction

Since the start of the HIV epidemic in the 1980’s formidable research efforts have been spent on the search for a protective vaccine that could reduce the numbers of new infections worldwide. As of now these efforts have been to no avail. The HIV-1 envelope glycoprotein (Env) trimer, located on the viral surface, is the sole target for neutralizing antibodies (NAbs), i.e. antibodies that prevent infection of new target cells. HIV-1 Env is highly diverse and antibodies need to cope with this diversity in order to become broadly active against different HIV-1 strains. Moreover, the Env trimer is heavily glycosylated to shield the underlying amino acids from recognition by the immune system. Despite these defenses, a proportion of HIV-1 infected individuals develop broadly neutralizing antibodies (bNAbs) that have evolved to accommodate glycans ^1^.

The Env trimer consist of three monomers of non-covalently linked gp120 and gp41, forming a heterotrimeric glycoprotein. These weak, non-covalent interactions have complicated the generation of stable Env trimers as vaccine antigens. However, advances in the last decade have led to the development of stable, soluble and native-like Env trimers, some of which are currently being evaluated in first-in-human studies ^2–7^. The prototype native-like Env trimer, BG505 SOSIP, contains a number of modifications to trap the soluble Env trimer in the pre-fusion state, allowing the presentation of quaternary bNAb epitopes, while occluding most non-neutralizing epitopes ^2^. Native-like Env trimer immunogens of various designs elicited NAb responses against the autologous virus, as well as sporadically against heterologous primary (tier 2) viruses in animals ^8–10^.

Despite success in eliciting autologous NAb responses against tier 2 viruses, the major challenge is to induce bNAb responses against conserved epitopes on the Env trimer. One site of interest is the CD4 binding-site (CD4bs), which is relatively conserved by virtue of the fact that nearly all HIV-1 viruses require CD4 binding for entry ^11^. HIV-1 infected individuals frequently develop bNAbs that target this conserved epitope ^12,13^. VRC01, isolated from an individual infected with a subtype B isolate, typifies the VRC01-class of CD4bs bNAbs ^13,14^. Many more class members have been described since, for example CH103 and CH235 ^15–18^. Both antibodies were isolated from an African individual that was infected with a subtype C strain ^16,17^. The CH103 lineage co-evolved with the transmitter/founder (T/F) virus CH505 and cooperated with the CH235 bNAb lineage ^17,18^.

Native-like Env trimers rarely induce CD4bs responses, but they can be modified to enhance the accessibility of the CD4bs and recruit bNAb precursors ^19,20^. This so-called germline-targeting trimer approach was in part fueled by an increasing understanding of the glycosylation of Env trimers and its impact on immunogenicity. It has been shown, that holes in the glycan shield represent immunodominant off-target epitopes, hence reducing their presence might further improve immunogenicity of conserved epitopes ^21–24^. T/F viruses with an intact glycan shield elicit bNAbs more effectively in humans ^25^. These findings suggest, that a recombinant native-like Env trimer with an intact glycan shield might be superior to an Env trimer with glycan holes at inducing NAbs.

Here we describe an immunization study in rabbits with a native-like trimer, AMC016 SOSIP.v4.2, derived from a subtype B Env, that has an intact glycan shield. A NAb response against heterologous tier 2 viruses of subtype B and a CD4bs-directed antibody specificity was induced in one animal. Additional immunizations with a combination of three different subtype B trimers further increased the neutralization breadth. The results should guide efforts to increase the consistency bNAb responses targeting the CD4bs.

## Results

### The AMC016 trimer induces NAbs against heterologous tier 2 viruses in one animal

Recently, we described a new trimer, AMC016 SOSIP.v4.2, that has an intact glycan shield as judged by the presence of PNGS at all conserved sites, as well as high glycan occupancy of these PNGS ^46^. The Env trimer is based on an *env* sequence of a subtype B virus, isolated 9 months post-seroconversion from an individual enrolled in the Amsterdam Cohort Studies for HIV/AIDS and who was classified as non-neutralizer ^26^. The AMC016 trimer induced no or only low-level autologous NAbs in rabbits, consistent with its dense glycan shield ^46^. These findings made the AMC016 trimer a suitable candidate for testing the hypothesis that Env trimers with an intact glycan shield might induce heterologous NAbs more efficiently than trimers with holes in the glycan shield such as the prototype BG505 SOSIP trimer ^21,25^.

In a previous study, rabbits (n=5) were immunized three times with the AMC016 SOSIP.v4.2 trimer, in combination with the adjuvant ISCOMATRIX ^46^. As it was previously shown that a fourth immunization can increase binding and neutralization titers against Env trimers, we continued the study by immunizing the five rabbits with the AMC016 trimer for a fourth time at week 36 (**Fig. 1A**) ^10^. The week 38 sera were tested in neutralization assays against a panel of viruses (**Fig. 1B** and **Table S1**). The autologous NAb response against the AMC016 virus was modest (median ID_50_ of 50) and 74- and 21-fold lower compared to the autologous NAb response induced by the BG505 and B41 SOSIP trimers after three immunizations, respectively (median ID_50_ of 3707 and 1048, respectively; n=10 for BG505 and n=5 for B41) ^27^. The BG505 and B41 trimers both lack conserved PNGS, resulting in autologous NAb responses that are dominated by specificities that target the holes in the glycan shield, explaining why they readily induce autologous NAb responses ^21,28,29^. Tier 1 subtype B viruses were poorly neutralized (median ID_50_ of 73, 57 and 20 for SF162, AMC008 and AMC011, respectively) (**Table S1**), while the murine leukemia virus (MLV), which functioned as a negative control, was not neutralized by any of the week 38 sera.

**Fig. 1.**
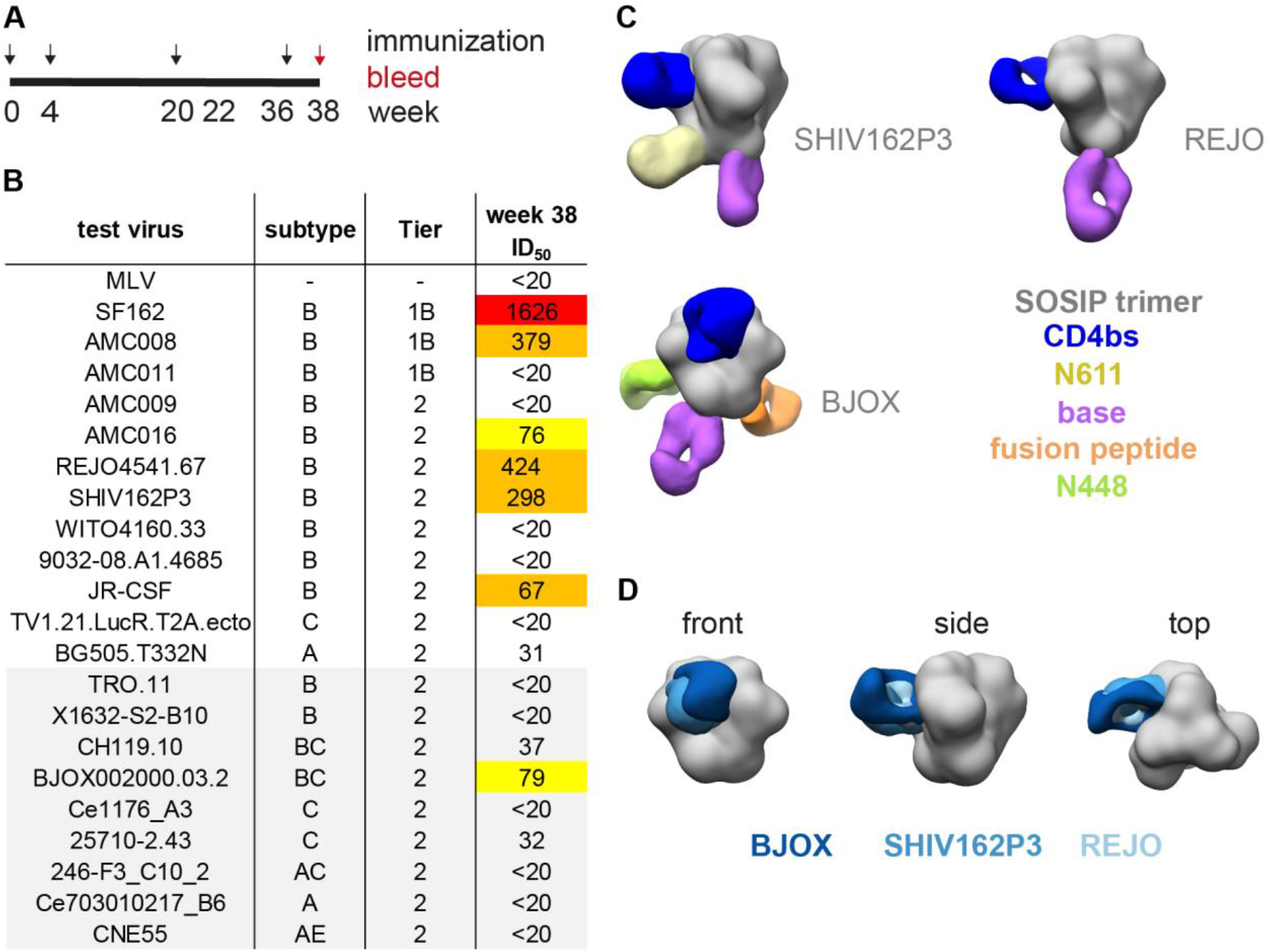
Immune response of rabbit UA0075 after immunization with the AMC016 trimer. (A) Immunization scheme. Indicated are the immunizations with AMC016 SOSIP.v4.2 (black arrows) and the bleed at week 38 (red arrow). (B) Neutralization titer (ID_50_) at week 38 against the autologous virus and heterologous tier 1 and tier 2 viruses. Viruses of the 9-virus global panel are highlighted in grey. The ID_50_ values for the five immunized rabbits can be found in Table S1. White: no neutralization, yellow: weak neutralization, orange: moderate neutralization, red: strong neutralization. (C) Fabs of week 38 polyclonal IgG were complexed with trimers based on the neutralized heterologous tier 2 viruses. The complexes were analyzed via negative stain EM. The targeted epitopes are indicated. (D) Overlay of the week 38 CD4bs responses (different shades of blue), targeting the three trimers of panel C.

A panel of 17 heterologous tier 2 viruses were tested, including the 9-virus global panel, that reflects the diversity of circulating viral strains from subtype A, B, C and recombinant viruses ^30,31^. One animal, rabbit UA0075, neutralized some of these viruses with an ID_50_ ≥40 (ID_50_ of 424, 298, 67 and 79 for REJO, SHIV162P3, JR-CSF and BJOX, respectively) (**Fig. 1B** and **Table S1**). The neutralization breadth, defined as the percentage of viruses neutralized with an ID_50_ ≥40 or with an ID_50_ ≥100, was 24% and 12%, respectively. When assessing the neutralization breadth within subtype B by only considering the 10 heterologous tier 2 viruses from subtype B, the serum from animal UA0075 neutralized 40% of the subtype B viruses with an ID_50_ ≥40 and 20% with an ID_50_ ≥100. Overall, while the AMC016 trimer induced a weak autologous NAb response in rabbits after four immunizations, the serum of animal UA0075 neutralized a subset of heterologous tier 2 viruses with moderate potency.

### The AMC016 trimer induces a cross-reactive CD4bs antibody response in animal UA0075

We analyzed the heterologous antibody response in animal UA0075 in more detail. Using electron microscopy-based polyclonal epitope mapping (EMPEM), we investigated whether we could identify a shared epitope responsible for the neutralization of heterologous tier 2 viruses ^32^. Polyclonal serum IgG was purified from the serum of rabbit UA0075 and digested with Papain to obtain Fabs (see Material and Methods). The Fabs were complexed with the autologous AMC016 trimer and trimers derived from the heterologous tier 2 viruses REJO, SHIV162P3 and BJOX that were neutralized by the serum (Brinkkemper et al., manuscript in preparation) ^10^. The complexes were then analyzed via negative-stain electron microscopy (NS-EM) (**Fig. 1C**) ^32^. It was not possible to obtain 3D reconstructions with the AMC016 trimer, as the trimer dissociated after being complexed with the Fabs, a phenomenon that has been previous observed with EMPEM analyses of sera from animals immunized with native-like trimers ^33^. However, we successfully obtained reconstructions with the three heterologous trimers. A non-neutralizing epitope on the trimer base (purple) was targeted on the REJO, SHIV162P3 and BJOX trimers. This is consistent with previous findings that the exposed base of native-like soluble trimers is immunodominant ^32^. The reconstruction of the SHIV162P3 trimer also revealed an antibody specificity bound to the N611 region (yellow), which is also commonly seen because the N611 PNGS is often underoccupied creating a hole in the glycan shield ^34^. The fusion peptide (orange) and N448 region (neon green) were targeted on the BJOX trimer. The most remarkable finding was an antibody specificity against the CD4bs (blue) that recognized all three trimers (**Fig. 1C**), with a very similar angle of approach (**Fig. 1D**). Native-like trimers rarely elicit CD4bs specificities unless deliberately modified to do so ^19,20^. Thus, immunization with the AMC016 trimer resulted in a CD4bs antibody response in animal UA0075. The antibody specificity cross-reacted with three Env trimers that corresponded to the neutralized heterologous tier 2 viruses.

### A trivalent subtype B trimer combination broadens the NAb response in animal UA0075

We previously reported that rabbits immunized with a combination of trimers induced sporadic tier 2 NAb more readily than animals immunized with a monovalent trimer ^10^. We posed that trimers which are genetically not too distinct, e.g., from the same subtype, might facilitate the induction of a NAb response to shared epitopes. In order to boost the already existing heterologous NAb response in animal UA0075, we elected to further immunize this animal with a combination of different subtype B trimers. We hypothesized that it would be beneficial if the immunogens represent the viral diversity within subtype B, to increase the neutralization breadth of viruses from the same subtype. To find sequences that fulfill these requirements, subtype B *env* sequences were obtained from the Los Alamos Database on HIV/AIDS and used for a maximum-likelihood analysis (http://www.hiv.lanl.gov/) (**Fig. 2A**; Material and Methods). The sequences were subsampled to represent viral diversity in time (1985 – 2019) and location (Africa, Asia, Europe, North America, South America and Oceania). We selected three *env* sequences, which are from different geographical locations: B41 (New York, USA), AMC008 (Amsterdam, The Netherlands) and SFU (Canada) (**Fig. 2A**). The B41 and AMC008 (based on *env* from individual H18818) trimers were previously described, whereas the SFU trimer was newly generated for this study ^35–37^. In contrast to the AMC016 trimer, the B41, AMC008 and SFU proteins miss one or more conserved PNGS (N130 and N289 for B41, N234 for AMC008 and N234 for SFU). The B41 virus was not neutralized by week 38 serum of rabbit UA0075 (data not shown).

**Fig. 2.**
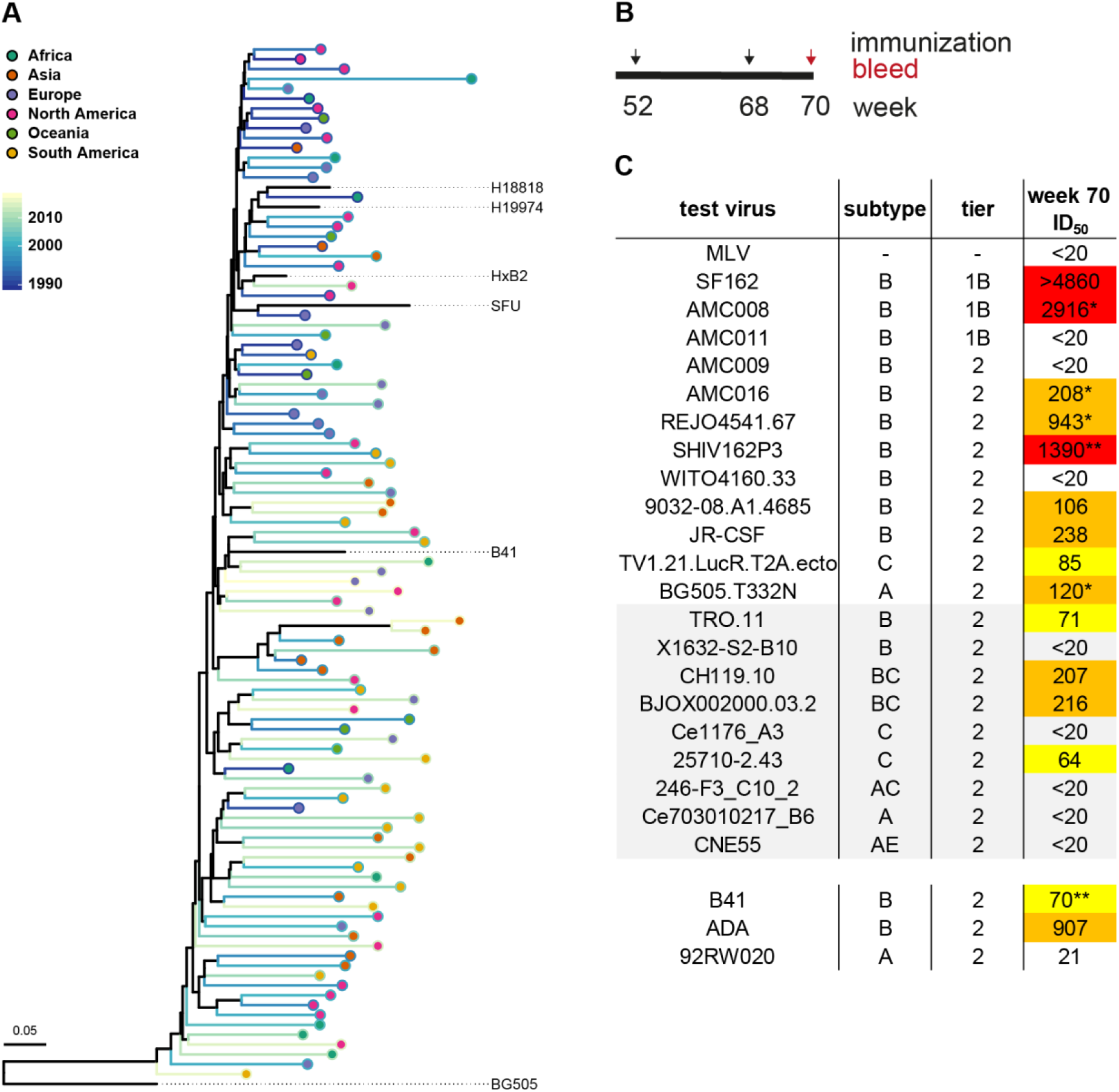
Immune response of rabbit UA0075 after additional immunizations with a trivalent subtype B combination. (A) Phylogenetic tree of subtype B viruses. Viral strains representing the diversity within subtype B in terms of time and place. Indicated are the three *env* sequences used for the trivalent combination: H18818 (source of AMC008 SOSIP), B41 and SFU, as well as H19974 (source of AMC016 SOSIP) ^35,36,46^. (B) Immunization scheme, continuation of Fig. 1A. Indicated are the two additional immunizations with the trivalent combination (black arrows) and the final bleed (red arrow). (C) ID_50_ neutralization titer at week 70. The same panel of viruses were tested as in Fig. 1B and the same color code applies. Viruses of the 9-virus global panel are highlighted in grey. Additional test viruses are listed below the table.

The SOSIP.v5.2 stabilizing mutations were introduced into the SFU sequence, the protein expressed in HEK293T cells and the supernatant analyzed in an enzyme-linked immunosorbent assay (ELISA) (**Fig. S1A**). BG505 SOSIP.v4.2 was taken along for comparison. The binding of bNAbs 2G12 and VRC01 to SFU and BG505 was comparable. The quaternary structure-preferring bNAbs PGT145, PGT151, PG9 and the gp120-gp41 interface bNAb 35O22 bound to both proteins, albeit with lower affinity to SFU. The SFU SOSIP protein was then expressed in HEK293F cells and affinity purified using PGT151. The purified SFU SOSIP.v5.2 protein was trimeric as assessed by BN-PAGE (**Fig. S1B**) and fully cleaved between the gp120 and gp41 subunits as assessed by SDS-PAGE (**Fig. S1C**). An analysis of the SFU trimer with NS-EM and hydrophilic interaction chromatography-ultra performance liquid chromatography (HILIC-UPLC) showed, that the trimer attained a native-like conformation (>95%) and was rich in oligomannose glycans (Man_5-9_: 56.2%, Man_9_: 24.4%) (Table 1), as expected from a native-like trimer ^2,38^.

**Table 1.** Biophysical properties of SFU SOSIP.v5.2

| Table 1 Biophysical properties of SFU SOSIP.v5.2 |  |  |  |
| --- | --- | --- | --- |
| Production |  | Yield (mg/L) | 2.96 <sup>a, b</sup> |
| Morphology | NS-EM | Native-like trimers (%) | 100 <sup>b</sup> |
| Glycan composition | HILIC-UPLC | Man<sub>5-9</sub> (%) | 56.2 <sup>b</sup> |
|  |  | Man<sub>9</sub> (%) | 24.4 <sup>b</sup> |
| <sup>a</sup>: Results were obtained from 293F cell-expressed and PGT151-purified SOSIP trimers |  |  |  |
| <sup>b</sup>: Results were obtained with D7324-tagged proteins |  |  |  |

The trivalent trimer combination, consisting of AMC008 SOSIP.v4.2, B41 SOSIP.v4.1 and SFU SOSIP.v5.2, was then used to immunize rabbit UA0075 intramuscularly with 22.5 μg of each trimer (amounting to a total of 67.5 μg trimer protein) at week 52 and 68 (**Fig. 2B**). The week 70 serum was tested in neutralization assays. Overall, the neutralization titers against tier 2 viruses increased from week 38 to week 70 (**Fig. 2C**). The autologous viruses AMC016, AMC008 and B41 were neutralized with an ID_50_ of 208, 2916 and 70, respectively (3- and 8-fold increase in ID_50_ for AMC016 and AMC008). The neutralization response against the autologous SFU virus was not determined, as no infectious virus could be produced (see Materials and Methods). The same heterologous tier 2 virus panel, as used for the analysis of week 38 sera, was then assessed (**Fig. 2C**). The NAb response against viruses that were already neutralized at week 38 increased. The subtype B and BC viruses REJO, SHIV162P3, JR-CSF, 9032-08.A1.4685, TRO.11, CH119 and BJOX were neutralized with an ID_50_ of 943, 1390, 238, 106, 71, 207 and 216, respectively. Similarly, the NAb response against the subtype A virus BG505 and the two subtype C viruses 25710 and TV1.2 increased (ID_50_ of 120, 64 and 85, respectively). The neutralization of REJO, SHIV162P3, JR-CSF, BG505, CH199, BJOX and 25710 increased by 2− to 6-fold (calculated for the viruses that were neutralized at week 38 with an ID_50_>20). In addition, the week 70 serum was tested against other heterologous tier 2 viruses. The subtype A virus 92RW020 was not neutralized, whereas the subtype B viruses ADA was neutralized with an ID_50_ of 907. The negative control virus, MLV, was not neutralized by serum of UA0075. The tier 1B virus SF162 was efficiently neutralized, whereas AMC011 was not (ID_50_ of 4860 and 20, respectively).

Overall, when using ID_50_ ≥40 as the cut-off the serum of rabbit UA0075 neutralized 59% of the viruses of the tested virus panel at week 70, representing an increase in breadth of 35%. Moreover, the serum neutralized 88% of the subtype B viruses tested (increase of 48%). When using a more stringent cut-off for neutralization (ID_50_ ≥100), the neutralization breadth in the week 70 serum was 41% for the entire panel and 75% for subtype B viruses (increase of 29% and 55%, respectively). We conclude that the trivalent subtype B combination boosted and broadened the NAb response within subtype B, but also broadened the response across other subtypes.

### The broad neutralization in the serum of animal UA0075 is IgG mediated

In order to validate the observation that the serum of animal UA0075 had broadly neutralizing activity, we studied neutralization of the IgG fraction in the serum. Due to limitations of week 70 serum, we used serum of the final bleed, at week 73, i.e. five weeks after the last immunization. First, we obtained data on the NAb response at week 73 and compared it to the week 70 NAb response (**Fig. 3A**). The neutralization titers waned substantially between week 70 and week 73, but broad neutralization was still observed. The ID_50_ values decreased 15-fold and 17-fold for SF162 and AMC008, respectively, 2-fold for AMC016, 4-fold for B41, REJO and SHIV163P3, and 6-fold for BG505. We next purified polyclonal IgG from the week 73 serum. The purified IgG was obtained in the same volume as the initial serum and used in a neutralization assay in the same dilutions as the serum (**Fig. 3B** and Materials and Methods). The neutralization titers of IgG were comparable to the serum, with a maximum difference of 2-fold for AMC008 and SHIV162P3. Both the serum and IgG did not neutralize MLV, B41 and BG505. We conclude that the broad neutralization in animal UA0075 was IgG mediated.

**Fig. 3.**
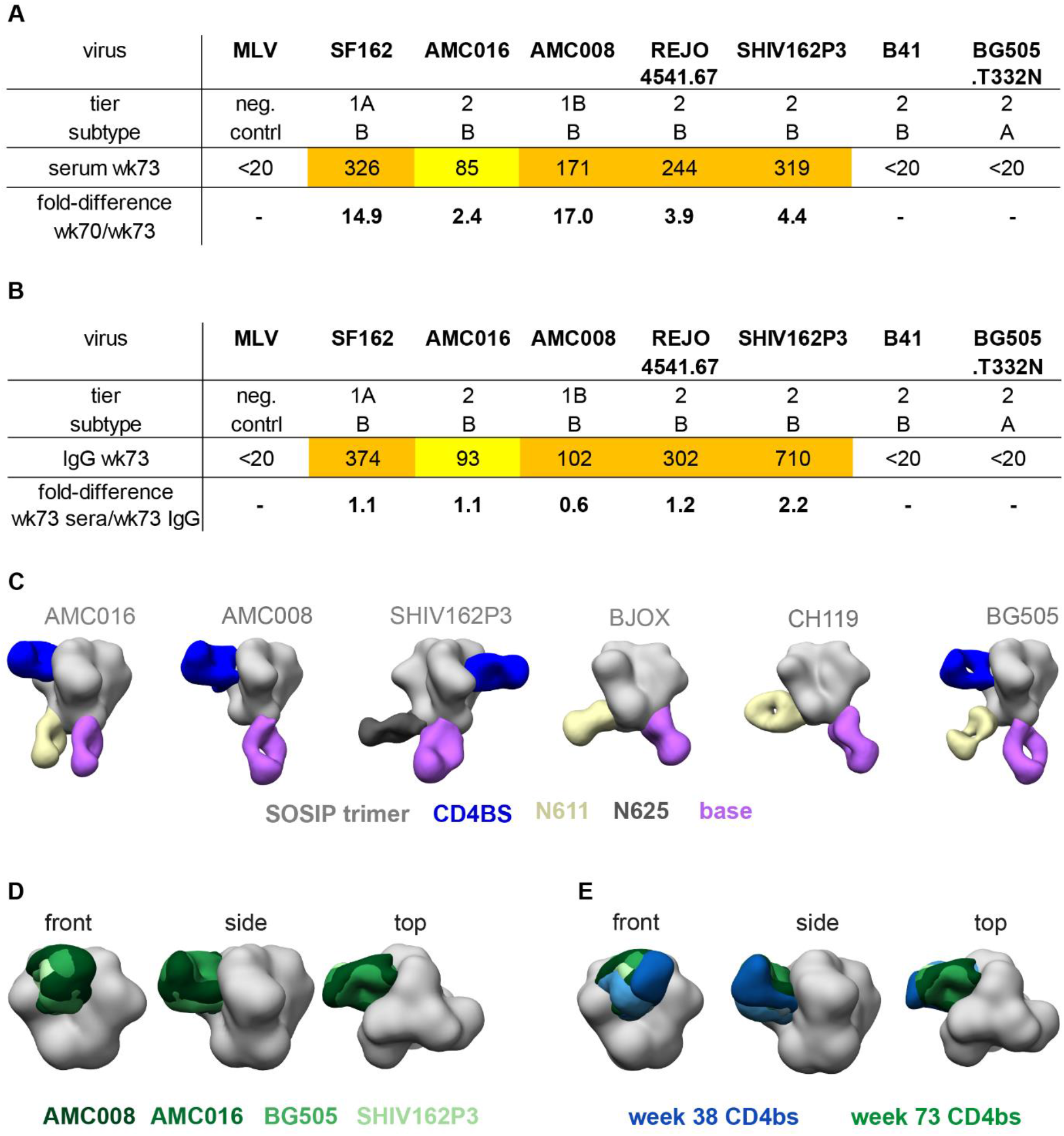
Characterization of the immune response of rabbit UA0075 at week 73. (A) ID_50_ titer at week 73 and comparison with the ID50 of week 70. (B) Comparison of the ID50 of serum and polyclonal IgG at week 73 (see Material and Methods for details). (A) and (B) The fold-difference is indicated for ID_50_>20. The same color code was used as in Fig. 1B. (C) EMPEM analysis of week 73 Fabs complexed with autologous and heterologous SOSIP trimers. The complexed trimers and the targeted epitopes are indicated. (D) Overlay of the CD4bs specificities of the AMC008, AMC016, SHIV162P3 and BG505 trimers. (E) Overlay of the CD4bs specificities at week 38 (blue) and week 73 (green).

### The cross-reactive CD4bs response persists after the trivalent booster immunizations

We were interested whether the trivalent subtype B booster combination affected the antibody specificity for the various epitopes on the autologous and heterologous trimers. The purified IgG from the week 73 serum was digested to obtain polyclonal Fabs. The Fabs were then complexed with the autologous AMC016 and AMC008 trimers and the heterologous SHIV162P3, BJOX, B41, REJO, CH119 and BG505 trimers (**Fig. 3C**). The B41 and REJO trimers dissociated into monomers and dimers when complexed with the serum Fabs and the target epitopes could not be assessed. While we had observed the same dissociation with the AMC016 trimer and the week 38 Fabs, it was possible to complex the AMC016 trimer with week 73 Fabs. Overall, the dominant specificities were the same as those observed at week 38. Serum specificities directed against the trimer base recognized all six trimers, while specificities directed against the N611 region and the CD4bs recognized four of the six trimers. Some changes were observed when comparing the reactivities of the week 38 and week 73 sera using the BJOX and SHIV162P3 trimers. The specificities against the N448 region, the fusion peptide and CD4bs were not detected anymore when assessed with the BJOX trimer, whereas an antibody targeting the N611 region was now detectable. For SHIV162P3, there was no binding of an antibody to the N611 glycan, but a new antibody specificity that bound to the N625 region.

While the Fabs directed to the trimer base and the N611 site represent non-NAb specificities, the broadly reactive CD4bs-directed Fabs may represent NAbs. An overlay of the CD4bs specificities that targeted the four trimers at week 73, showed that the antibodies have the same angle of approach and probably target the same region on the trimers (**Fig. 3D**). Similarities in the angle of approach were also observed for the CD4bs-specificities in the week 38 and the week 73 CD4bs samples (**Fig. 3F**; week 38 is indicated in blue; week 73 in green). Thus, animal UA0075 developed a cross-reactive CD4bs-directed response that remained present after the booster immunizations.

### The CD4bs specificity in animal UA0075 shows resemblance to bNAb CH235

To further specify the CD4bs epitope targeted in animal UA0075, the NS-EM structure of CD4bs specificity was overlayed with the bNAbs VRC01, CH103 and CH235.12 (**Fig. 4A**, **Fig. 4B** and **Fig. 4C**). The UA0075 CD4bs antibody bound to the CD4bs at a site somewhat closer to the trimer apex compared to VRC01 and with a different angle of approach (**Fig. 4A**; VRC01 in blue, BG505 in dark grey; PDB ID 6V8X) ^39^. In contrast, the UA0075 CD4bs antibody has a similar angle of approach as the CH103 antibody and they both target a similar region on the trimer (**Fig. 4B**; CH103 in red and BG505 gp120 in light grey; EMDB 6250) ^40^. Similarities in angle of approach and targeted epitope were also observed for the comparison with CH235.12 (**Fig. 4C**, pink; EMDB 8079).

**Fig. 4.**
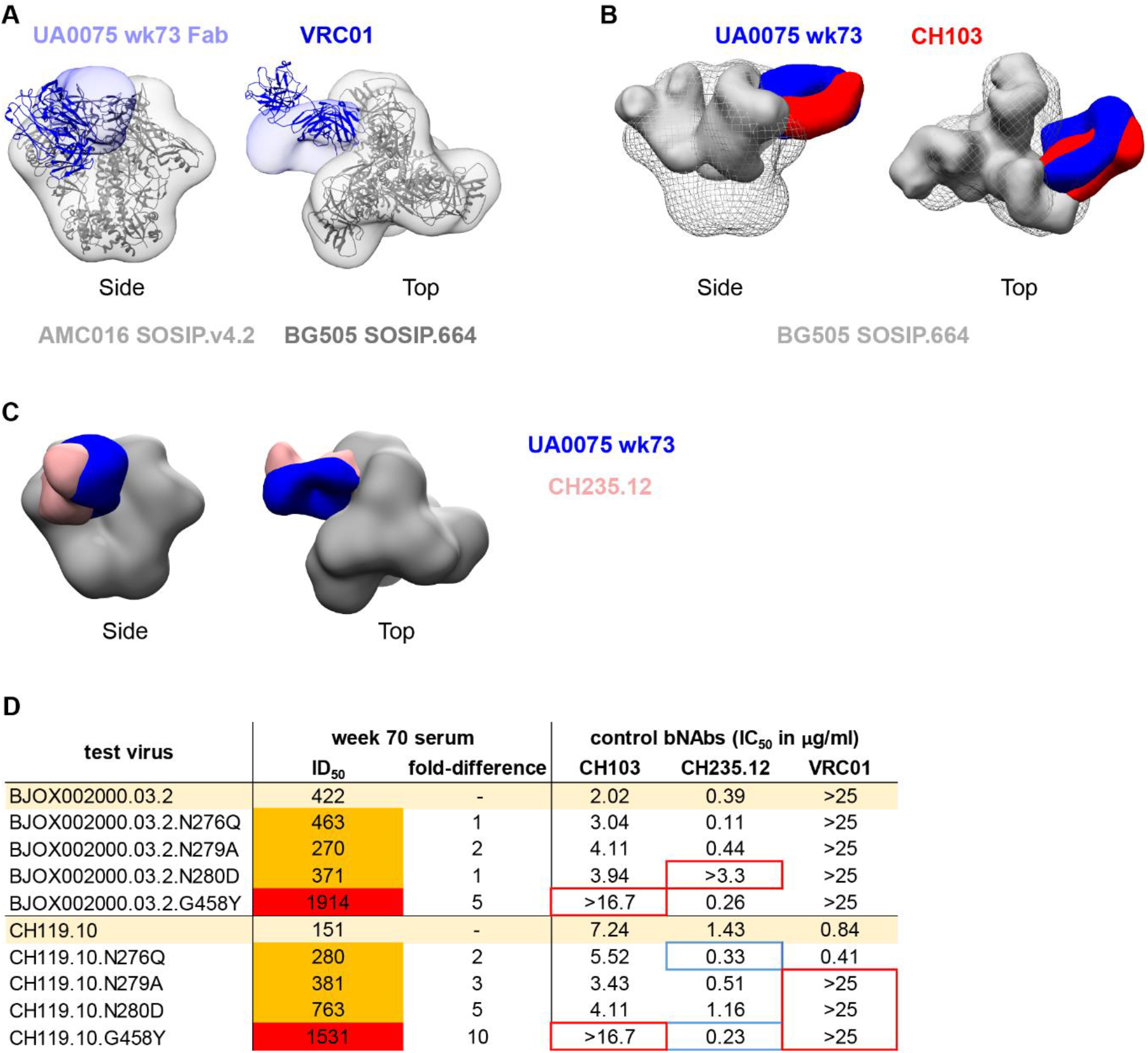
Characterization of the CD4bs antibody response of rabbit UA0075. (A) Overlay of the CD4bs antibody response of UA0075 (lilac), complexed with AMC016 SOSIP.4.2 (grey density), and the VRC01 (blue; PDB ID 6V8X) and BG505 SOSIP.664 structures (dark grey) ^39^. (B) Overlay of the CD4bs antibody of UA0075 (blue) with the low density structure of CH103 (red; EMDB 6250) complexed with BG505 SOSIP.664 (grey) ^40^. (C) Overlay of the UA0075 CD4bs specificity (blue) and CH23.12 (nude; EMDB 8079). (D) Neutralization titer (ID_50_) against BJOX and CH119 mapping viruses, which contain point mutations in the CD4bs. Indicated is the fold-difference in ID50 compared to the parental virus. The same color code was used for the ID_50_ values as in Fig. 1B. The bNAbs VRC01, CH103 and CH235.12 were taken along for comparison and the IC_50_ (in μg/ml) was calculated. Absence of neutralization is indicated by a red box, while a blue box indicates an increase in neutralization.

We next studied the UA0075 CD4bs specificity by using CD4bs mapping viruses based on the heterologous tier 2 viruses CH119 and BJOX. The mapping viruses contain single point mutations in the CD4bs that result in glycan deletions: N276Q, N279A, N280D and G458Y (**Fig. 4D**). The three bNAbs that share similarities with the UA0075 CD4bs specificity were tested against these mapping viruses. Neutralization of VRC01 was abrogated by N279A, N280D and G458Y, in the context of CH119. Neutralization of CH103 was eliminated by the G458Y mutation in both CH119 and BJOX. In contrast, N276Q and G458Y in the context of CH119 increased neutralization titers for CH235.12. We then tested the serum of UA0075 against the mapping viruses. Neutralization was not removed by any mutation and the mapping viruses were neutralized with a similar ID_50_ compared to the parental viruses (**Fig. 4D**). The exception was G458Y, that resulted in a 5- and 10-fold increase in ID_50_ (for BJOX and CH119, respectively). A weaker (2-fold) enhancement of neutralization was also observed with CH119 N276Q. In terms of neutralization, the UA0075 NAb response bears most similarities with the bNAb CH235.12.

To conclude, EM revealed that the CD4bs specificity in the serum of animal UA0075 shows resemblance to bNAbs CH103 and CH235.12 and the enhanced sensitivity of the G458Y point mutation in the BJOX and CH119 viruses was reminiscent of CH235.12.

## Discussion

In this manuscript we described the humoral immune response induced by a subtype B Env trimer, AMC016, followed by a trivalent subtype B combination. The study was initiated with the hypothesis that an Env trimer with an intact glycan shield might be able to induce NAbs. The autologous NAb response in all immunized animals was low, as expected, but one rabbit developed a cross-neutralizing antibody response against heterologous tier 2 viruses. Whether the intact glycan shield focused the immune response on a shared, conserved epitope, we cannot fully answer here, because a NAb response developed in inconsistently, i.e. only in one of five rabbits. One possible explanation for the inconsistency of the response is the presence of immunodominant, non-neutralizing epitopes, such as the trimer base or the underoccupied N611 PNGS, which creates a glycan hole ^32,34^. Presentation of the AMC016 trimer on nanoparticles combined with enhancing PNGS occupancy are among the strategies to improve the consistency of the NAb response induced by the AMC016 trimer ^3,4,41^.

We assume that the cross-reactive CD4bs specificity in animal UA0075 is responsible for the neutralization and also the neutralization breadth within subtype B. First, the other two dominant epitopes targeted, N611 and the trimer base, are non-neutralizing epitopes. Second, neutralization of heterologous tier 2 viruses was affected by mutations in the CD4bs. Further and detailed epitope mapping studies should strengthen the assumption that the CD4bs specificity is the dominant NAb specificity.

Interestingly, a rabbit CD4bs-directed mAb with neutralization breadth against heterologous tier 2 viruses was also described by Dubrovskaya et al ^42^. NFL trimer-liposomes were used in a prime-boost immunization regime. The priming Env trimer missed four glycans around the CD4bs, followed by subsequent immunizations with heterologous trimers with restored glycans from different subtypes. In both studies, a sporadic CD4bs response was elicited with an advanced prime-boost regime and carefully selected immunogens. Both CD4bs specificities differed from VRC01. While the prime NFL trimer-liposome was specifically designed to make the CD4bs accessible, the AMC016 trimer possessed a complete glycan shield.

A complementary approach to focus the immune response on conserved epitopes such as the CD4bs is represented by germline-targeting trimers ^6,19,43^. One well-studied germline-targeting trimer, GT1 serves as a prime immunogen to initiate CD4bs-directed NAb lineage. In addition to generating direct contacts with VRC01-class germline precursors, one feature of the GT1 design is the removal of glycans around the CD4bs to make the site accessible for germline antibodies, akin to the strategy described by Dubrovskaya et al. ^42^. After GT1 priming, the CD4bs-directed antibody responses would have to be ‘shaped’ and ‘polished’ by immunogens in which the glycans are being re-introduced, with the aim to guide the immune system towards the production of bNAbs that can accommodate these glycans. We propose that the AMC016 trimer, with its intact glycan shield and the potential to induce a CD4bs NAb response, might be a suitable trimer for the final ‘polishing’ immunization.

In the immunization regime used here, a trivalent combination of subtype B trimers were used to further broaden and boost the NAb response. Interestingly, all trimers of the trivalent combination had holes in the glycan shield, but the three major specificities that were induced after AMC016 trimer immunization, i.e. those against the CD4bs, the N611 region and the trimer base, were still present after immunizations with the trivalent combination. The choice of subtype B trimers for the trivalent combination might have facilitated the broadening of the CD4bs-directed NAb specificities, but we can speculate that filling the glycan holes of the components of the trivalent combination might further improve the CD4bs response.

We showed here, that the AMC016 SOSIP trimer with an intact glycan shield can induce a potent NAb response in rabbits, that efficiently neutralized heterologous tier 2 viruses of subtype B. The trimer induced a cross-reactive CD4bs-directed antibody response that could be boosted and broadened by a trivalent combination of subtype B trimers. The results should guide efforts to induce CD4bs-directed NAb responses more consistently.

## Acknowledgements

We thank the participants of Amsterdam Cohort Studies on HIV and AIDS for providing their samples. This work was funded by the U.S. National Institutes of Health grant P01 AI110657 (to ABW and RWS), and by the Bill and Melinda Gates Foundation (BMGF) grants OPP1132237 and INV-0020220 (to RWS) and OPP115782 (to ABW). ABW was supported by the IAVI Neutralizing Antibody Center through the Collaboration for AIDS Vaccine Discovery grant OPP1196345/INV-008813, funded by the BMGF. RWS received funding from the European Union’s Horizon 2020 research and innovation program under grant agreement no. 681137. RWS is a recipient of a Vici grant from the Netherlands Organization for Scientific Research (NWO). MJvG is a recipient of an AMC Fellowship and a Mathilde Krim Fellowship from the American Foundation for AIDS Research (amfAR) (109514-61-RKVA). The Amsterdam Cohort Studies on HIV infection and AIDS, a collaboration between the Public Health Service Amsterdam, the Amsterdam UMC of the University of Amsterdam, Medical Center Jan van Goyen and the HIV Focus Center of the DC-Clinics, are part of the Netherlands HIV Monitoring Foundation and financially supported by the Center for Infectious Disease Control of the Netherlands National Institute for Public Health and the Environment. The funders had no role in study design, data collection and analysis, decision to publish, or preparation of the manuscript.

## Material and Methods

### Construct design, protein expression and purification

The recently described AMC016 SOSIP trimer is based on *env* of a biological clone isolated 9 months post-seroconversion, derived from infected individual H19974, enrolled in the MSM (men having sex with men) component of the Amsterdam Cohort Studies on HIV/AIDS (ACS) ^26^. H19974 did not develop bNAbs ^26^. The AMC016 SOSIP.v4.2 trimer was described previously ^46^. BJOX SOSIP.v9, SHIV162P3 SOSIP.v9 and REJO SOSIP.v4.2 were used for the EMPEM analysis (Brinkkemper et al., manuscript in preparation; Bontjer et al., manuscript in preparation). H18818, the source of the AMC008 SOSIP.v4.2 trimer was also enrolled in the ACS, whereas the B41 *env* was obtained from an infected individual in New York and expressed as B41 SOSIP.v4.1 ^35,36^. The SFU *env* gene was obtained from a chronically infected individual living in Canada. The codon-optimized SOSIP.v5.2 SFU *env* was ordered at GenScript (Piscataway, NJ) and cloned into the pPPI4 expression vector, expressed in HEK-293T and HEK-293F cells and purified using the bNAb PGT151 (Bontjer et al., manuscript in preparation).

### Rabbit immunization

Immunizations were performed at Covance (Denver, PA, USA), under approval number C0048.15. New-Zealand rabbits were immunized intramuscularly with 40 μg of AMC016 SOSIP.v4.2 at week 0, 4, 20 and 36, followed by immunizations at week 36, 56, 70 with 67.5 μg of the trivalent trimer combination, consisting of AMC008 SOSIP.v4.2, B41 SOSIP.v4.1 and SFU SOSIP.v9 (22.5 μg for each component) ^46^. ISCOMATRIX (CSL Ltd., Parkville, VIC, Australia) was used as adjuvant for all immunizations. Blood was drawn two weeks after each immunization and the week 38, 70 and 73 sera used for neutralization assays, IgG purification and EMPEM analysis.

### Neutralization assay

A TZM-bl cell neutralization assay was performed, as previously described, to analyze the NAb response at weeks 38, 70 and 73 ^8,44^. Polyclonal IgG from week 73 was purified to compare the serum neutralization vs IgG neutralization. The IgG was obtained in the same volume as the serum for purification. In a neutralization assay, the IgG and serum were treated exactly the same in terms of input and dilution. The gene for the SFU virus was ordered as a pseudovirus, using the original *env* sequence. The SFU pseudovirus was not infectious. The BJOX and CH119 mapping viruses were designed and tested at Duke University.

### HILIC-UPLC for N-glycan profiling

To obtained information on the glycan composition of the SFU SOSIP.v9 (see Table 1), the trimer was analyzed with HILIC-UPLC, as previously described ^38^

### NS-EM

The SFU SOSIP.v9 trimer was analyzed with NS-EM to acquire information on the native-like confirmation (see Table 1). The method was previously described in detail ^45^.

### EMPEM analysis

Serum IgG from UA0075 at week 38 and week 73 was purified using a Prot A column and digested with Papain to obtain Fabs ^10,32^. Fabs were complexed with SOSIP trimers, purified via size-exclusion, analyzed with NS-EM and the images processed ^32^.

### Phylogenetic tree of HIV-1 subtype B strains

HIV-1 envelope sequences of subtype B were obtained from the Los Alamos HIV Database (http://www.hiv.lanl.gov/). To represent overall diversity, the sequences were randomly subsampled such that at least one sequence was included each year (1985 – 2019) for each continental region (Africa, Asia, Europe, North America, Oceania and South America). Additional subtype B envelope sequences were added to the sequences: B41, SFU, H19974 (AMC016) and H18818 (AMC008) ^35,36^. The subtype B sequence HxB2 was included as a reference for numbering and the subtype A sequence BG505 for rooting the tree ^2^.

