## Supplementary Material for "CD4 binding-site antibodies induced by a subtype B HIV-1 envelope trimer"

**Table S1** Immunogenicity of AMC016 SOSIP.v4.2 immunized rabbits

| | | | **Immunogen** | **AMC016 SOSIP.v4.2** | | | | |
|---|---|---|---|---|---|---|---|---|
| | | | **week** | **38** | | | | |
| **test virus** | **subtype** | **tier** | **Rabbit ID/ institute** | **UA0072** | **UA0073** | **UA0074** | **UA0075** | **UA0076** |
| **MLV** | - | neg. ctrl. | AMC | <20 | <20 | <20 | <20 | <20 |
| | | | DUMC | <20 | <20 | <20 | <20 | <20 |
| **SF162** | B | 1A | AMC | 69 | 2587 | 73 | 1626 | 32 |
| **AMC008** | B | 1B | AMC | 57 | 300 | 22 | 379 | <20 |
| | | | DUMC | 24 | 24 | <20 | 178 | <20 |
| **AMC011** | B | 1B | AMC | <20 | <20 | <20 | <20 | <20 |
| | | | DUMC | <20 | <20 | <20 | <20 | <20 |
| **AMC016** | B | 2 | AMC | <20 | 50 | 1016 | 76 | <20 |
| | | | DUMC | <20 | 26 | 441 | 71 | <20 |
| **AMC009** | B | 2 | AMC | <20 | <20 | <20 | 21 | <20 |
| **REJO4541.67** | B | 2 | AMC | <20 | 27 | <20 | 424 | <20 |
| | | | DUMC | <20 | <20 | <20 | 214 | <20 |
| **SHIV162P3** | B | 2 | AMC | <20 | 21 | 33 | 298 | <20 |
| **WITO4160.33** | B | 2 | AMC | <20 | <20 | 34 | <20 | <20 |
| | | | DUMC | <20 | <20 | <20 | <20 | <20 |
| **B41** | B | 2 | AMC | <20 | <20 | <20 | <20 | <20 |
| **BG505.T332N** | A | 2 | AMC | <20 | <20 | <20 | 31 | <20 |
| **TRO.11** | B | 2 | DUMC | <20 | <20 | <20 | <20 | <20 |
| **X1632-S2-B10** | B | 2 | DUMC | <20 | <20 | <20 | <20 | <20 |
| **CH119.10** | BC | 2 | DUMC | <20 | <20 | <20 | 37 | <20 |
| **BJOX002000.03.2** | BC | 2 | DUMC | <20 | <20 | <20 | 79 | <20 |
| **Ce1176_A3** | C | 2 | DUMC | 26 | <20 | <20 | <20 | <20 |
| **25710-2.43** | C | 2 | DUMC | <20 | <20 | <20 | 32 | <20 |
| **246-F3_C10_2** | AC | 2 | DUMC | <20 | <20 | <20 | <20 | <20 |
| **Ce703010217_B6** | A | 2 | DUMC | <20 | <20 | <20 | <20 | <20 |
| **CNE55** | AE | 2 | DUMC | <20 | <20 | <20 | <20 | <20 |

**Table S1** Neutralization titers of rabbits immunized with the AMC016 trimer at week 38. Serum of the five rabbits were tested against the autologous virus and heterologous viruses of tier 1 and tier 2. Indicated are the serum dilutions that reduce infection by 50% in a TZM-bl cell assay. White: No or very weak neutralization ($ID_{50}$ <20 and $ID_{50}$ 20-40, respectively). Yellow: Weak neutralization ($ID_{50}$ 40-100). Orange: Moderate neutralization ($ID_{50}$ 100-1000). Red: Strong neutralization ($ID_{50}$ >1000). Neutralization of the autologous and heterologous tier 2 viruses were repeated for verification and the $ID_{50}$ values were averaged. Neutralization assays were performed at Academic Medical Center (AMC). Data, obtained at Duke University Medical Center (DUMC), is shown for comparison.

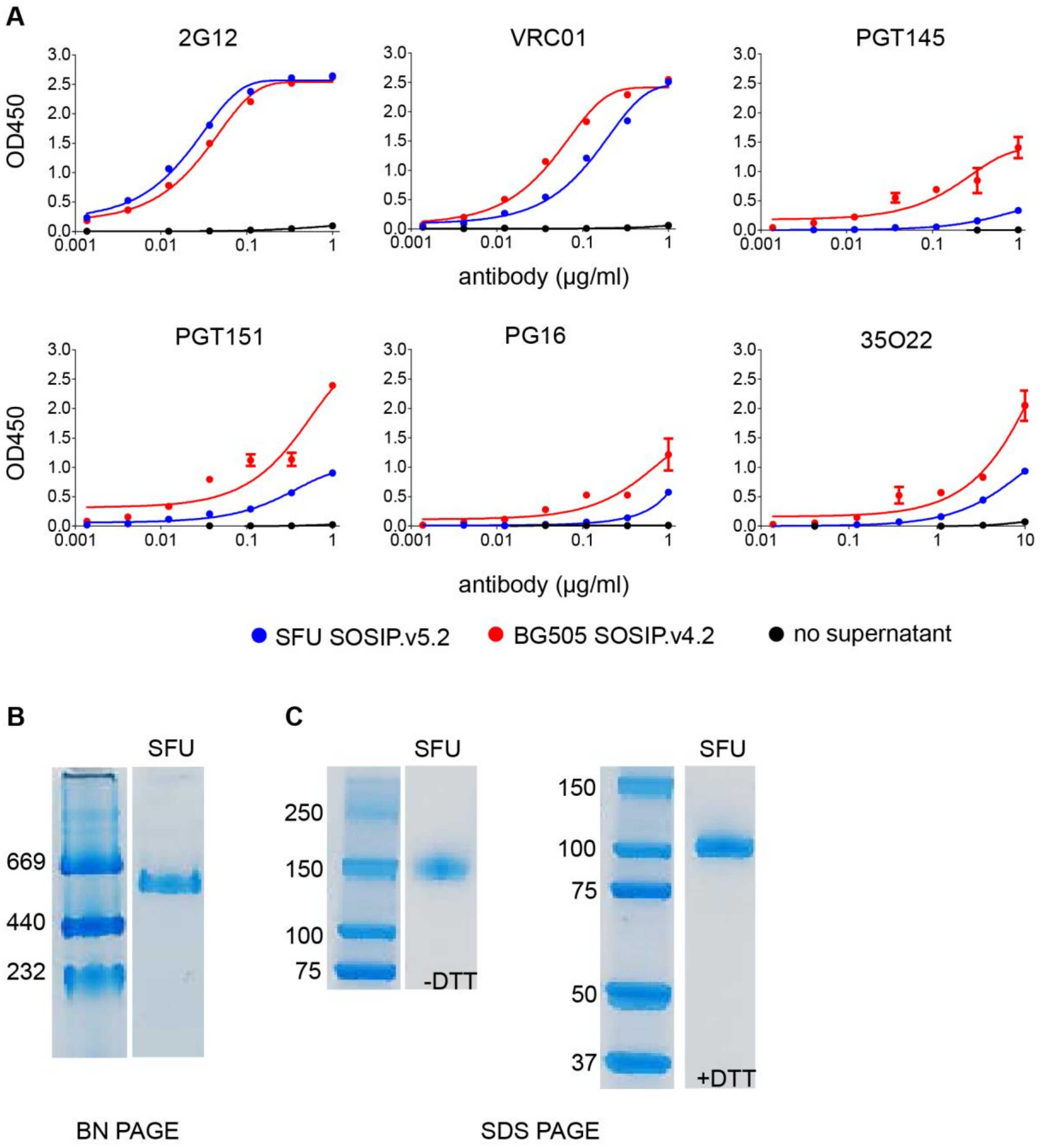


**Fig. S1** Biophysical and biochemical characterization of SFU SOSIP.v5.2. (A) ELISA using supernatant of SFU transfected 293T cells. BG505 SOSIP was taken along for comparison. As a negative control, no supernatant was used in the assay. (B) BN-PAGE with PGT151 purified SFU trimer. (C) SDS-PAGE with purified SFU trimer under non-reduced (-DTT) and reduced (+DTT) conditions.
